# A collection of genetically engineered *Populus* trees reveals wood biomass traits that predict glucose yield from enzymatic hydrolysis

**DOI:** 10.1101/124396

**Authors:** Sacha Escamez, Madhavi Latha Gandla, Marta Derba-Maceluch, Sven-Olof Lundqvist, Ewa J. Mellerowicz, Leif J. Jönsson, Hannele Tuominen

## Abstract

Wood represents a promising source of lignocellulosic biomass for the production of bio-based renewables, especially biofuels. However, woody feedstocks must be improved to become competitive against petroleum. We created a collection of *Populus* trees consisting of 40 genetically engineered lines to modify and to better understand wood biomass properties. A total of 65 traits were measured in these trees and in the corresponding wild-type clone, including growth parameters, wood anatomical and structural properties, cell wall composition and analytical saccharification. The relationships between saccharification of glucose and biomass traits were investigated using multivariate data analysis methods and mathematical modeling. To circumvent potential trade-offs between biomass production and saccharification potential, we also estimated the “total-wood glucose yield” (TWG) expected after pretreatment and 72h of enzymatic hydrolysis from whole trees. A mathematical model estimated TWG from a subset of 22 wood biomass traits with good predictivity (Q^2^ = 0.8), while saccharification of glucose could be predicted from seven biomass traits (Q^2^ = 0.49). Among the seven diagnostic traits for saccharification, four also affected biomass production, such as the ratio of S- to G-lignin which was beneficial for saccharification but detrimental for growth. The contents of various matrix polysaccharides appeared important for predicting both saccharification and TWG, including low abundance monosaccharides. In particular, fucose and mannose contents negatively correlated with TWG, apparently by negatively associating with biomass production. Both biomass production and saccharification, and hence TWG, negatively correlated with arabinose and rhamnose contents, suggesting that these low abundance monosaccharides represent markers/targets for improving feedstocks.

## Significance statement

Lignocellulosic biomass from trees represents a source of sugars to produce renewable commodities instead of petroleum-based products. However, both the production processes and feedstocks need to be improved for large-scale implementation. This study provides information and tools for the selection or the engineering of trees for use as renewable feedstocks. We generated 40 genetically engineered *Populus* lines and characterized their biomass properties in relation to the estimated sugar yield after enzymatic hydrolysis. Mathematical modeling was used to predict sugar yield based on wood traits and to identify diagnostic traits that represent tentative tools for selecting superior trees.

## Introduction

Sugars extracted from wood biomass represent a promising source of renewable biofuels and other green chemicals to sustainably replace petroleum-based products (1-4). In particular, the biochemical conversion of lignocellulosic biomass holds great potential (3), although improvements are needed at every step of the process (3), starting with the feedstocks.

Tree species from the *Populus* genus represent interesting lignocellulosic feedstocks because they exhibit rapid growth even on marginal lands and are widely and efficiently cultivated (5,6). Furthermore the genomes of several *Populus* species have been sequenced (5, 6). Research efforts have focused on improving the biomass production of Populus feedstock (7-10). However, for biochemical conversion it is important to also consider woody biomass recalcitrance hindering the release of sugars from the wood, which therefore requires harsh pretreatments, synonymous of higher costs (11).

Biomass recalcitrance has been studied in natural variants of the *Populus* genus (12-14), showing that lignin amount and composition affected saccharification (14), and revealing parts of the genetic relationships underlying lignin as well as other biomass traits (12, 13). Parallel approaches relied on targeted genetic engineering of xylem cell walls, resulting in trees less recalcitrant to enzymatic saccharification, although sometimes at the expense of growth (15-22). Interestingly, saccharification or sugar conversion could be improved by altering the composition of matrix polysaccharides (16, 17), reducing the amount of lignin (18) or modifying lignin composition (19, 20). Together, these studies provide useful information for future breeding or genetic engineering programs as well as potential feedstocks. However, translating these tools and knowledge into practice requires further research into aspects such as trade-off between reduced recalcitrance and increased biomass production.

The present study contributes to bridging this knowledge gap by characterizing the relationship between biomass traits and susceptibility to enzymatic saccharification in a population of transgenic hybrid aspen (*Populus tremula × tremuloides*; hereafter *Populus*) known as the BioImprove collection. We estimated the glucose yield after pretreatment and 72h enzymatic hydrolysis from the total wood biomass of each tree to identify diagnostic traits for the creation and selection of not only less recalcitrant but overall superior trees with increased sugar yield. Such selection could be applied in current breeding programs to enhance biochemical conversion rates. Furthermore, our collection of transgenic trees theoretically comprises combinations of traits that are not currently found in nature, paving the way for a deeper biological understanding of woody biomass and of the ways to improve it.

## Results

### The BioImprove *Populus* collection provides a trait library for characterizing wood biomass properties and glucose yield

We investigated the relationships between wood traits and the potential of woody biomass for enzymatic saccharification in *Populus* trees by altering the expression of genes putatively regulating wood biomass properties. Thus, the expression of 39 genes was targeted in a collection of 40 transgenic *Populus* lines (Dataset S1). These lines, as well as the wild-type T89 clone, were analyzed for 3 growth-related traits, 20 cell wall chemistry traits, 20 wood anatomy and structural traits and 22 saccharification traits (Dataset S2), thus generating a broad wood-related trait library. Notably, a wide variation was observed for major growth traits such as height and diameter (Fig 1a,b), for traits critical for biomass recalcitrance such as lignin content and lignin monomer composition (Fig 1c,d) and for analytical saccharification traits such as glucose release after 72h of enzymatic hydrolysis without or after pretreatment (Fig 1e,f). This variation in quantitative traits between lines is valuable as it allows us to decipher how wood properties influence traits of interest, such as glucose yield.

**Figure 1:**
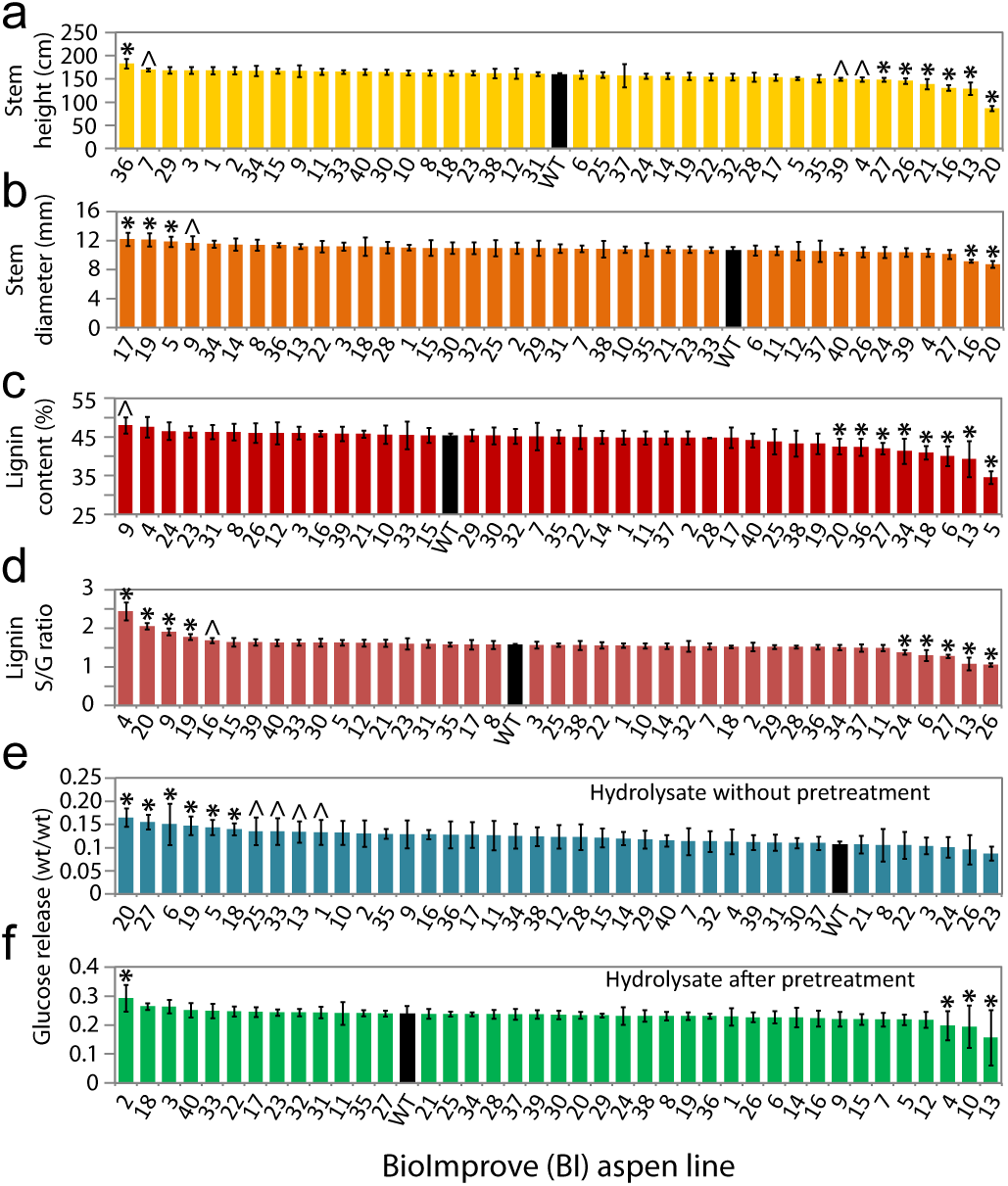
The BioImprove poplar collection provides a wide variation in major traits. (a,b) Growth-related traits: stem height (a) and stem diameter (b). (c,d) Biomass recalcitrance-related traits: relative lignin content within the detected pyrolysate from biomass (a) and ratio of S- to G-units within the lignin polymer (b). (e,f) Saccharification-related traits: glucose release after a 72h enzymatic hydrolysis without (e) or after (f) pretreatment. Each histogram represents the average value for a transgenic line (color) or wild-type (black). Error bars represent standard deviation. * and ^∧^ indicate statistically significant differences from wild-type (p<0,05 and p<0,1 respectively) following a post-ANOVA Fisher’s test (n=3-5).

Notably, saccharification is usually expressed as the relative amount of sugar released per unit of biomass, which reflects the recalcitrance rather than the yield of an entire tree. Trees with high saccharification may concomitantly suffer from growth defects, which may nullify the *in fine* yield of these trees. Instead, ideal trees for biochemical conversion of biomass should combine high saccharification with sufficient growth to ensure superior yield from their total wood biomass. To reflect this, we created a combinatorial trait – a tree’s “total-wood glucose yield” (TWG; Fig 2a), as glucose is the most prominent product of saccharification, with diverse uses. The TWG was used for ranking the *Populus* trees based on how much glucose could be released from their whole stems after pretreatment and 72h enzymatic hydrolysis. Interestingly, several BioImprove *Populus* lines exhibited significantly different TWG compared with the wild-type trees (Fig 2b).

**Figure 2:**
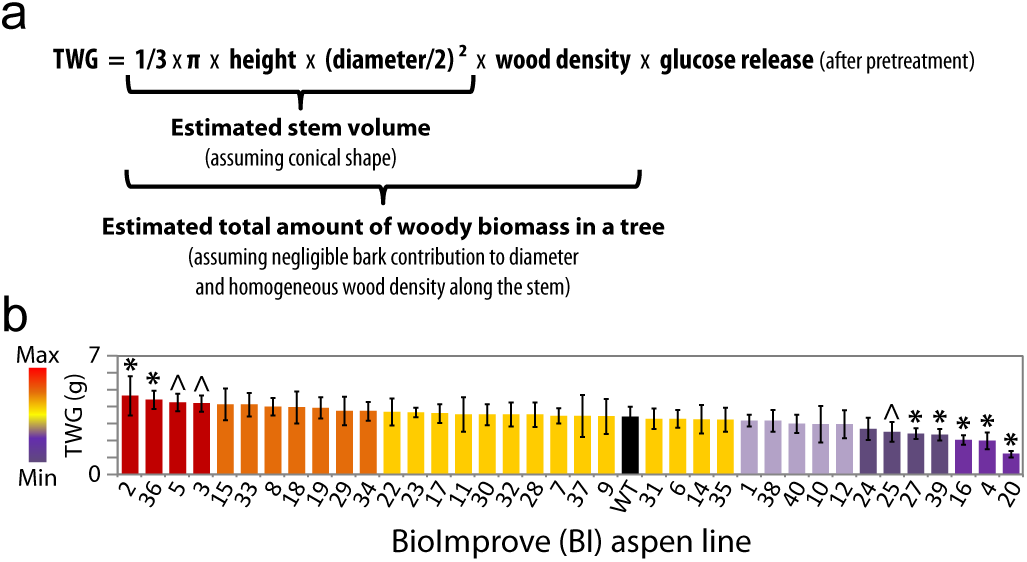
The BioImprove lines display different potential for total wood glucose yield (TWG) (a) Formula for estimation of a tree’s total-wood glucose yield after pretreatment and 72h enzymatic hydrolysis, assuming conical shape, negligible bark contribution to diameter and homogeneous wood density. (b) TWG of the BioImprove poplar lines (in g). Each histogram represents the average value for a transgenic **Populus** line (color) or wild-type (black). Error bars represent standard deviation. * and ^∧^ indicate statiscally significant differences from wild-type (p<0,05 and p<0,1 respectively) following a post-ANOVA Fisher’s test (n=3-5).

To identify variation in traits that could separate the lines based on TWG, we first performed a principal component analysis (PCA). The resultant PCA model displayed nine significant principal components (PCs; Dataset S3) explaining 78.6% of the variation in the data: 30.2% were explained by the two first PCs. Neither of the two first PCs (Fig S1) nor any other combination of PCs could separate the lines based on their TWG. Hence, TWG varied differently between lines than the traits underlying the main biological variation, implying the need for a different method to identify biomass properties associated with TWG.

### Certain traits are associated with total wood glucose yield

To overcome the limitations of the PCA analysis, we compared the 38 *Populus* lines whose TWG could be calculated (Fig 2b) using a supervised, predictive multivariate analysis. Orthogonal projection of latent structures (OPLS; (23)) enables us to distinguish the variation related to a variable of interest, for instance TWG, from the unrelated (orthogonal) systematic variation. An OPLS model was generated which could separate the *Populus* lines with respect to TWG (Fig 3a) in a significantly predictive manner (Q^2^ = 0.75).

**Figure 3:**
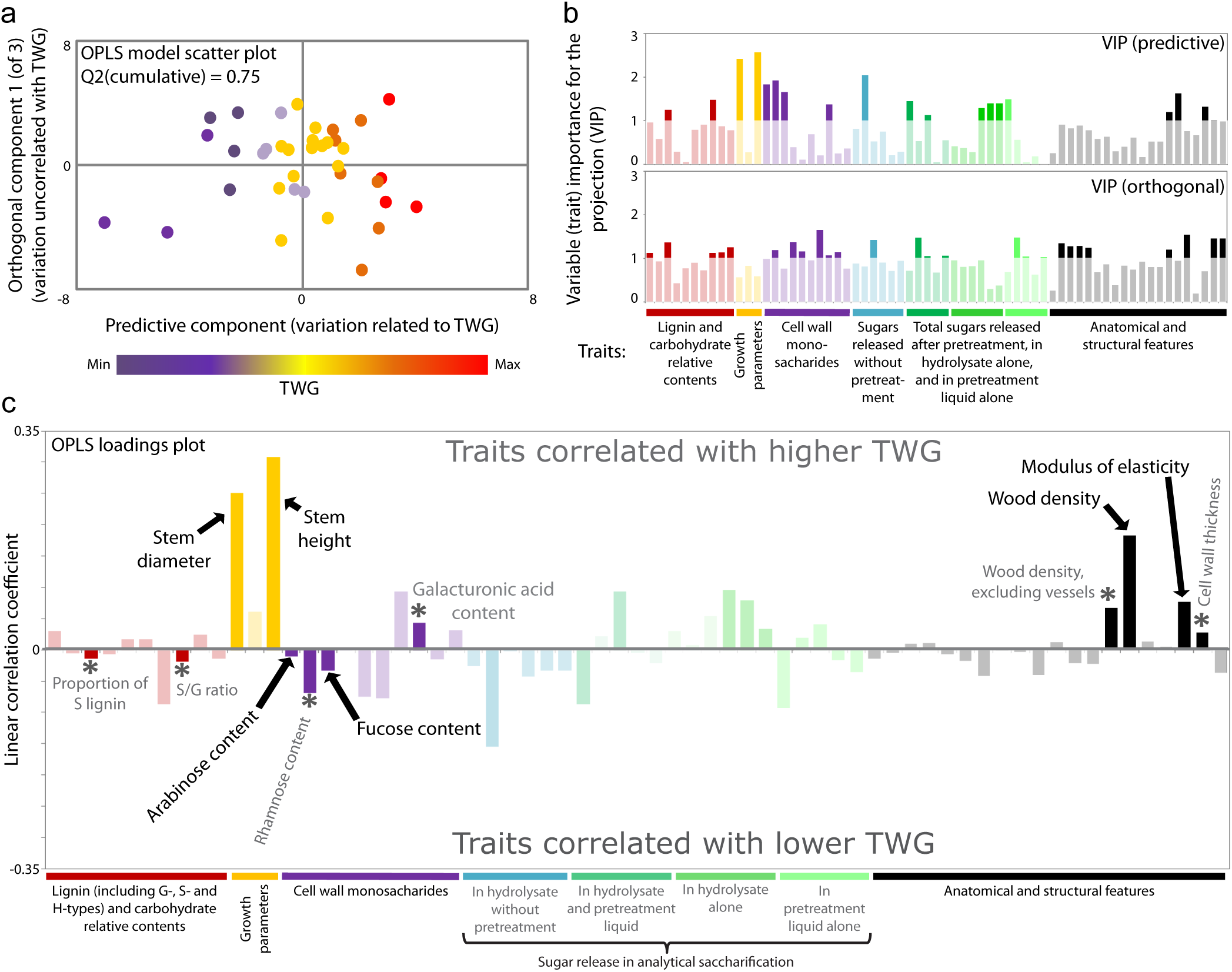
Certain traits contribute more than others to predicting TWG. (a) OPLS scatter plot showing the separation of *Populus* lines (dots) horizontally along the predictive component for TWG. Vertical separation indicates variation not correlated with TWG. The lines were coloured by TWG. (b) Plots showing the variable importance for the projection (VIP) value for each trait for the predictive part of the model (up) and for the orthogonal part of the model (down). VIP values over 1 indicate important traits. (c) Contribution of each trait to the OPLS model. Appart from saccharification traits, traits with a VIP value over 1 for the predictive part of the model were emphasized by black text and arrows. Traits marked by (*) and annoted in grey are important (VIP value over 1) for both the predictive and the orthogonal part of the model. Q2 scores over 0.5 indicate significant predictivity of a model.

In order to identify the traits which contribute most to predicting TWG in the BioImprove collection, we calculated each trait’s VIP (variable importance for the projection) values for both the TWG-predictive and TWG-orthogonal parts of the OPLS model (Fig 3b). Attempts to use VIP in order to reduce the number of traits used to predict TWG also strongly reduced the model’s predictivity. Although the model relied on all 65 traits, VIP values (Fig 3b) indicated that some traits contributed more to predicting TWG than others. We therefore relied on VIP to focus our interpretation on the traits which appeared more important for TWG prediction (Fig 3c).

Among the traits related to biomass production, wood chemistry and wood structure and anatomy, 12 traits were significantly associated with TWG in the OPLS model (Fig 3c). Height, diameter and wood density were positively associated with TWG (Fig 3c), as expected from the fact that TWG is a composite trait which integrates these measurements. Consistent with the contribution of density to TWG, increased wood stiffness (modulus of elasticity) and cell wall thickness were also associated with higher TWG (Fig 3c). Interestingly, galacturonic acid content was positively associated with TWG while arabinose, rhamnose and fucose contents were negatively associated with TWG (Fig 3c), showing that quantitatively minor cell wall compounds could influence TWG. Increases in S-type lignin content and in the ratio of S- to G-type lignin were weakly but significantly negatively associated with TWG in the OPLS model.

### Mathematical modeling predicts TWG from a subset of wood biomass traits

The OPLS analysis revealed the possibility of predicting TWG from wood biomass traits in our dataset. However, our OPLS model relies on all traits, making it informative but difficult to apply to predict TWG from future datasets. Hence, we attempted to generate a model to predict TWG from only a subset of wood biomass traits and which could also be used with future datasets to verify the general applicability of this model.

First, separate models were generated for each of the four traits, height, diameter, wood density and glucose release after pretreatment, from which TWG is calculated (Dataset S4). Then, by replacing each term in the TWG equation (Fig 2a) with the corresponding model, we obtained a composite model to predict TWG (Dataset S4). In that way, the potential effect of predictive traits on TWG could be traced down to an effect on saccharification, on biomass production, or even both. The resulting composite model could predict TWG (Fig 4) with significant accuracy (Q^2^ = 0.80). In contrast to the OPLS model, which included all the traits in the dataset, our composite mathematical model relied solely on twenty-two biomass traits (Table 1). Any attempt to reduce this number of traits also greatly reduced the model’s predictivity, suggesting that TWG is a complex trait emerging from intricate biological interactions.

**Figure 4:**
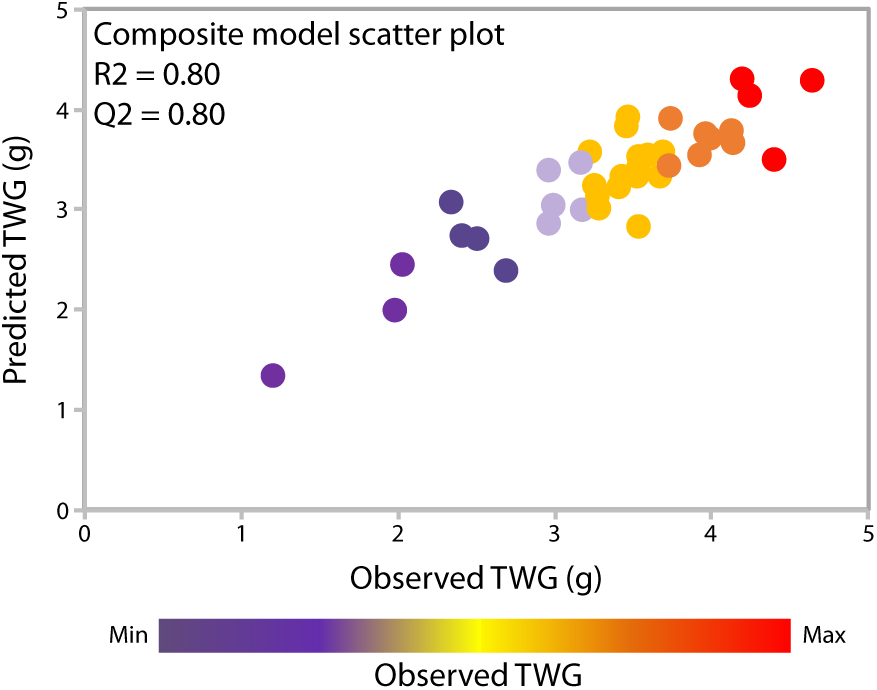
TWG can be predicted by a specific subset of traits in a composite model. Scatter plot showing for each poplar line (dots) the observed TWG (x-axis) versus the predicted TWG (y-axis). This plot displays the results of a composite model which is distinct from the OPLS model. Q2 scores over 0.5 indicate significant predictivity of a model.

**Table 1:** Wood traits contributing to predicting TWG in the composite model.

| Traits (contributing to either of the models) | Direction of contribution of a trait to predicting TWG in the composite model |
| --- | --- |
| Proportion of S lignin | Positive |
| Ratio of S-type to G-type lignin | Negative* |
| Arabinose content | Negative* |
| Rhamnose content | Negative* |
| Fucose content | Negative |
| Modulus of elasticity (stiffness) | Positive * |
| Cell wall thickness | Positive * |
| Xylose content | Negative |
| Mannose Content | Negative |
| 4-O-methylglucuronic acid content | Negative |
| Galactose content | Positive |
| Extractable glucose content (non crystalline) | Negative |
| Proportion of G lignin | Positive |
| Proportion of H lignin | Positive |
| Proportion of non-annotated phenolic coumpounds | Positive |
| Proportion of overall lignin | Positive |
| Ratio of cell wall carbohydrates to lignin | Positive |
| Fraction of wood (cross-sectional) area occupied by fibers | Positive |
| Average (cross-sectional) longest radial width of fibers | Negative |
| Average (cross-sectional) longest tangential width of fibers | Negative* |
| Average number of fibers per wood area | Negative |
| Average cross-sectional area of fibers | Negative |

\* The direction of the contribution of this trait to TWG is based on the wild-type level because this trait’s relationship with TWG is overall non-monotonic (i.e. the direction is not constant).

The use of four individual models to construct the composite model enables predicting the four individual variables which compose TWG. While stem height, diameter and wood density are easily measured traits and therefore do not need to be predicted from wood biomass traits, it remains informative for future feedstocks improvement strategies to know how wood anatomical and chemical traits influence not only saccharification but also biomass production. In addition, one of the individual models allows for predicting glucose release after pretreatment, a difficult trait to measure, based on only seven traits and with acceptable accuracy (Q^2^ = 0.49; Dataset S4). As expected (14), this model found the ratio of S- to G-type lignin to positively correlate with saccharification, but to negatively correlate with stem diameter (Dataset S4). Three other traits that predicted glucose release were also associated with stem height and/or diameter (Dataset S4), demonstrating the interplay between biomass recalcitrance and biomass production. Of particular interest, low abundance cell wall monosaccharides such as rhamnose and arabinose were associated with both glucose release and biomass production (Dataset S4). While rhamnose was negatively associated with both biomass production and saccharification, and therefore with TWG (Table 1), arabinose had a non-linear relationship to saccharification and a negative impact on biomass production (Dataset S4), also resulting in a mainly negative impact on TWG (Table 1), consistent with the OPLS model. Hence, our modeling approach revealed quantitatively minor matrix polysaccharides as targets for selection or engineering of woody biomass in *Populus*.

## Discussion

Our study identified putative diagnostic wood traits for the selection of trees with overall enhanced glucose yield. Previous studies had started unravelling the links between saccharification and other wood properties by studying populations of natural variants (12-14). Our work provides new, additional information despite relying on a smaller population of younger trees thanks to a different approach. We measured numerous traits from transgenic lines, which allowed us to analyze biological replicates, and to generate combinations of traits which may not occur in nature. Furthermore, the estimated TWG enabled us to circumvent potential trade-offs between biomass production and recalcitrance. Examples exist in the literature of (genetically modified) trees with improved saccharification (18, 19) which is offset by a concomitant growth reduction (19) or counter-acted by defects in xylem hydraulics (18, 21, 22). Consequently, the use of the TWG calculation or of similar proxies that integrate biomass production and sugar release, in addition to traditionally monitored saccharification, may help future studies to identify superior trees.

Lignin content and composition are considered major determinants of biomass recalcitrance to saccharification, as verified in a large population of undomesticated *Populus trichocarpa* in which the S- to G-lignin ratio was positively correlated with glucose release after pretreatment (14). Consistently, in our model for glucose release after pretreatment the S- to G-lignin ratio was a positive contributor (Dataset S4), confirming the relationship between lignin composition and biomass recalcitrance. However, when considering TWG, which integrates biomass production and saccharification, both our OPLS model and composite model revealed a negative impact of the S- to G-lignin ratio (Fig 3c, Table 1), likely because of its detrimental effect on stem diameter (Dataset S4). This observation interrogates the usefulness of increased S- to G-lignin ratio to improve the overall sugar yields in biochemical conversion of feedstocks.

In an earlier study, lignin content negatively correlated with saccharification in *Populus* trees with a ratio of S- to G- lignin below 2 (14), a range within which nearly all our trees fell (Fig 1d). Unexpectedly, lignin content did not negatively correlate with glucose release after pretreatment in our PCA analysis (Fig S1; Dataset S3) or in pairwise comparison (Spearman’s rank correlation rs = -0.09). Furthermore, lignin content did not contribute to predicting glucose release after pretreatment in the corresponding model (Dataset S4), indicating that lignin content did not greatly contribute to recalcitrance in our trees. Such discrepancy between studies in the effect of lignin content on saccharification may be explained by differences in methods, age of the trees, genetic background, degree of domestication, and/or growth environment. Indeed, analysis of a set of *Populus trichocarpa* trees grown at two locations revealed different degrees of negative correlation between lignin content and glucose release depending on growing site (12). On the other hand, these negative correlations between lignin content and saccharification were never statistically significant (12). In addition, Studer et al. (14) noted that a number of trees did not follow the general correlations between lignin content or composition and saccharification, leading them to propose that factors other than lignin can greatly influence biomass recalcitrance. Hence, the above observations are consistent with the emerging view that the woody biomass recalcitrance to saccharification is more complex than previously thought (for review see (11)), and that lignin does not necessarily affect sugar release.

An important source of variation between our lines may have been associated with tension wood (Fig S1). Tension wood is regarded as a determinant of wood recalcitrance because it has been found to improve saccharification in willow, although at the expense of biomass production (24). In the *Populus* genus, tension wood is associated with changes in cell wall monosaccharide composition such as decreases in xylose and mannose contents and concomitant increases in rhamnose, galacturonic acid and galactose contents (25, 26). Monosaccharide contents were also associated with TWG in our models (Table 1). The negative association of TWG with xylose and mannose contents together with the positive association of galactose content with TWG (Table 1) are consistent with an overall beneficial role of tension wood on TWG. However, the negative associations of rhamnose, non-crystalline glucose and arabinose contents with TWG (Fig 3c, Table 1) cannot be explained by tension wood, suggesting that differences in pectin and hemicelluloses composition unrelated to tension wood also influence TWG. This observation is in line with studies in *Arabidopsis thaliana* (27, 28) and *Populus* (15, 29) suggesting hemicelluloses as a promising target for biotechnological engineering of biomass to increase saccharification without growth penalty.

It is interesting to note that among the matrix polysaccharides significantly associated with TWG (Fig 3c, Table 1, Dataset S4), fucose, mannose, rhamnose and arabinose constitute quantitatively modest components of the wood biomass. Neither mannose nor fucose contributed to predicting saccharification but they negatively correlated with stem diameter and stem height, respectively (Dataset S4). Consequently, the composite model identified mannose as a negative contributor to TWG (Table 1) while fucose negatively correlated with TWG in both the composite model and the OPLS model (Fig 3c, Table 1). Arabinose and rhamnose were associated with both saccharification and biomass production in the individual models constituting our composite model (Dataset S4) such that they had an overall negative association with TWG in both the composite model (Table 1) and the OPLS model (Fig 3c). Hence, lower arabinose and rhamnose contents represent markers for increased biomass production combined with lower recalcitrance, either for the engineering or the selection of superior feedstocks. These results exemplify the importance of measuring quantitatively modest traits in future studies.

Our work relies on the use of transgenic lines designed to target specific genes, which allows us to discuss the potential genetic basis for the observed phenotypes. For instance, we found four *Populus* lines displaying significantly (p < 0.1) higher TWG than the wild type. While the causal link between the targeted genes and the improved TWG will require further investigation, three out of these four genes (in BI-2, BI-3 and BI-36) have not yet been characterized in relation to wood formation. This suggests that there probably remains a wealth of uncharacterized candidate genes which may provide markers for the selection of superior *Populus* trees or which represent targets for the biotechnological improvement of growth and biomass properties.

In conclusion, we uncovered a set of putative diagnostic traits for a combination of improved growth and biomass properties for saccharification which provides tentative tools for selecting *Populus* genotypes with high TWG. Indeed, *Populus* trees have been subject to domestication for a long time and there consequently exist numerous breeding populations (30-32) from which promising individuals could be selected.

## Materials and Methods

More detailed descriptions of the Materials and Methods are available in the Supplementary Information.

### Plant material and growth conditions

To create the BioImprove collection, transgenic hybrid aspen (*Populus tremula × tremuloides* Michx.) T89 clones were derived partially from a gene mining program performed at SweTree Technologies AB and partially from individual research groups at Umeå Plant Science Centre. The genes and the types of transgenic modifications are described in Dataset S1. Trees were grown for two months in previously described greenhouse conditions (33). Each tree’s height, diameter and mean internode length were measured and 8 cm-long sections of wood were harvested 20 cm above ground to perform most analyses.

### Cell wall compositional analyses and saccharification

Relative contents of cell wall lignin and carbohydrates, as well as lignin composition, were measured by pyrolysis-gas chromatography/mass spectrometry and the data were processed as previously described (34).

Cell wall monosaccharides were extracted by methanolysis with 2M HCl/MeOH, derivatized by trimethylsilyl and measured as previously described (17).

Pretreatment and analytical saccharification by enzymatic hydrolysis were performed according to the same study (17), but using half the amount of enzyme mixtures for hydrolysis.

### Wood anatomical and structural features

SilviScan (CSIRO, Australia) measurements conducted at INNVENTIA were performed incrementally along a parallelepipedic radial piece of wood as described previously (35), followed by normalization to reflect the total cross-sectional area that it represents in the wood.

### Statistics, data analysis and modeling

Average trait values of all the lines were compared by ANOVA. The lines were compared pairwise by post-ANOVA Fisher’s tests while Spearman’s rank correlations allowed the comparison of traits across lines, both using Minitab 17 (Cleverbridge AG, Germany).

The PCA and the corresponding post-PCA OPLS (25) analyses were performed on all lines and all but 3 lines (BI-13, 21 and 26), respectively, using SIMCA 14.1 (MKS Data Analytics Solutions, Sweden). In the OLPS, traits related to saccharification were disregarded in our interpretation of TWG prediction because the effort intensive process of measuring saccharification, in addition to other traits, would allow direct calculation of TWG.

The composite model to predict TWG was composed of four individual models generated to predict stem height, stem diameter, wood density and glucose release after pretreatment. These models were selected based on their predictivity (Q^2^) out of numerous (≥30) models generated using R.

## Acknowledgements

The authors thank Lars Olsson, Laura Stefana Ganea Koyin, Junko Takahashi-Schmidt, Veronica Bourquin and Lorenz Gerber for helping with measuring traits. We thank the SweTree Technologies Research Team in Umeå, and especially Magnus Hertzberg, for producing and selecting many of the transgenic plants used in the study. We also thank Konrad Abramowicz and Åke Brannstrom for their strong input on mathematical modeling. We thank Tomas Skotare and Bastian Schiffthaler for their advice on data analysis

The work was funded by the Swedish Research Council Formas (232-2009-1698 to H.T.; 942-2015-84 to H.T.) and by the Swedish Foundation for Strategic Research (RBP14-0011 to E.M.). We also thank the Umeå Plant Science Centre Berzelii Centre in Forest Biotechnology, funded by the Swedish Research Council VR and the Swedish Governmental Agency for Innovation Systems, and the UPSC plant cell wall laboratory, supported by Bio4Energy and TC4F.

## Author contributions

HT originally designed the study with assistance from EJM, LJJ and SOL. MLG, SOL, SE and HT contributed to the experiments and data acquisition. SE analyzed the data with help from HT, SOL, LJJ, MLG, EJM and MDM. SE performed the mathematical modeling. SE and HT wrote the manuscript, with assistance from SOL, EJM, LJJ, MLG and MDM.

